# Demographic Modeling of Admixed Latin American Populations from Whole Genomes

**DOI:** 10.1101/2023.03.06.531060

**Authors:** Santiago G. Medina-Muñoz, Diego Ortega-Del Vecchyo, Luis Pablo Cruz-Hervert, Leticia Ferreyra-Reyes, Lourdes García-García, Andrés Moreno-Estrada, Aaron P. Ragsdale

## Abstract

Demographic models of Latin American populations often fail to fully capture their complex evolutionary history, which has been shaped by both recent admixture and deeper-in-time demographic events. To address this gap, we used high-coverage whole genome data from Indigenous American ancestries in present-day Mexico and existing genomes from across Latin America to infer multiple demographic models that capture the impact of different timescales on genetic diversity. Our approach, which combines analyses of allele frequencies and ancestry tract length distributions, represents a significant improvement over current models in predicting patterns of genetic variation in admixed Latin American populations. We jointly modeled the contribution of European, African, East Asian, and Indigenous American ancestries into present-day Latin American populations to capture the historical demographic events that have shaped genetic variation. Our inferred demographic histories are consistent across different genomic regions and annotations, suggesting that our inferences are robust to the potential effects of linked selection. In conjunction with published distributions of fitness effects for new nonsynonymous mutations in humans, we show in large-scale simulations that our models recover important features of both neutral and deleterious variation. By providing a more realistic framework for understanding the evolutionary history of Latin American populations, our models can help address the historical under-representation of admixed groups in genomics research, and can be a valuable resource for future studies of populations with complex admixture and demographic histories.

## Introduction

Genetic diversity among human populations has been shaped by recurrent periods of migration and admixture^1^. Relatively recent admixture across Latin America has resulted in a rich and complex history, shaped by the mixing of ancestries from multiple regions, including Indigenous American, European, and African populations. Each of these populations has its own unique demographic history, both prior to and following their arrival to the continent^2–4^. As a result, the genetic composition of Latin American populations depends on both the shared broad-scale history of global human expansion and distinct regional admixture events, leading to patterns of genetic diversity that can vary markedly among Latin American populations and between individuals. However, there is currently a lack of historical models that accurately capture the demographic history and genetic composition of Latin American populations.

Accurate demographic models are essential for investigating a range of evolutionary, epidemiological, and ecological questions, as well as for simulating realistic genetic data^5, 6^. But despite the importance of understanding the demographic dynamics of Latin American populations, current models often fail to fully capture the heterogeneity of their genetic diversity. One limitation is that such models do not incorporate the diverse ancestries that have contributed to the genetic makeup of Latin American populations. For example, previous studies have used East Asian populations as a proxy for Indigenous American ancestries^7, 8^, despite the genetic divergence between present-day Indigenous American and East Asian individuals due to many thousands of years of isolation and serial founder effects during the peopling of the Americas^9–12^.

Current models are also limited by simplistic admixture histories, which do not accurately reflect the complexity of admixture in Latin American populations^8, 13^. Admixture histories in the Americas vary across geographical regions^4, 14^, and Latin American populations have experienced admixture events that involved additional populations beyond those typically considered in demographic models. For example, admixture events in Mexico have also involved individuals who descended from East and Southeast Asian populations^15^. This highlights the need for more nuanced models to fully capture the diverse ancestral contributions to the genetic makeup of present-day populations.

The demographic histories of Latin American populations comprise both recent and deeper-in-time processes that shape patterns of genetic diversity^16–18^. Inferring demographic models for these cohorts is challenging, as it requires considering multiple historical timescales. At the most recent timescale, admixture between source populations with different present-day genetic ancestries has resulted in a mosaic of genetic ancestry in Latin America. In turn, variation within contributed ancestries has been shaped by the demographic histories of each source population. This deeper-in-time history therefore also contributes to genetic diversity among Latin Americans^19^. To fully capture the intricate histories of these populations, it is therefore essential to consider both of these timescales jointly in demographic inference.

In this study, we infer more comprehensive demographic models for multiple cohorts in Latin America that better capture the complex and multifaceted history of these populations. To do this, we employed high-coverage whole genome sequences from worldwide populations and from across Latin America, including Colombia, Mexico, Peru, and Puerto Rico. We also included 50 recently sequenced individuals of Indigenous ancestries from Mexico, which provide increased resolution to distinguish East Asian and Indigenous American ancestries^3, 20^. We integrated demographic inferences at different timescales using the distributions of allele frequencies within and between populations and the numbers and lengths of ancestry tracts within individuals. Using extensive genetic simulations, we show that the resulting models accurately reflect multiple features of genetic variation observed in Latin American cohorts.

Our inferred demographic models can be coupled with models of selection or quantitative traits to explore the joint effects of demography and selection in shaping genomic variation. We demonstrate this application by recovering patterns of variation among missense and nonsense mutations in protein-coding genes. This approach can be extended to model the genetic basis of disease susceptibility and other traits in Latin American cohorts, and should serve as a valuable resource for researchers studying evolutionary and medical genomics in Latin American cohorts and other populations with a recent genetic mixture.

## Results

### Inferring the deep demographic history through allele frequencies

We used the joint site-frequency spectrum (jSFS) to infer a demographic model for the broad-scale human expansions that resulted in populations representing sources of ancestry of recent admixture in Latin America. The jSFS is the distribution of allele frequencies across multiple populations and can be used to infer demographic parameters such as effective population sizes, divergence times, and migration rates^7^. Allele frequencies were computed in high-coverage whole genome data from two DNA sequencing projects: the 1000 Genomes Project (1KGP)^21, 22^ and the Mexican Biobank Project (MXB)^3, 20^. Specifically, we used data from the Yoruba in Ibadan, Nigeria (YRI, as a proxy for African ancestries, AFR), the Iberian population in Spain, (IBS, as a proxy for European ancestries, EUR), the Han Chinese in Beijing, China (CHB, as a proxy for East Asian ancestries, EAS), and the Indigenous American population in Mexico (as a proxy for American ancestries, AME) (Figure 1). These or related populations contributed to the genetic makeup of present-day Latin Americans^19^ and by using the jSFS, we aimed to identify major demographic events that have influenced their genetic diversity.

**Figure 1.**
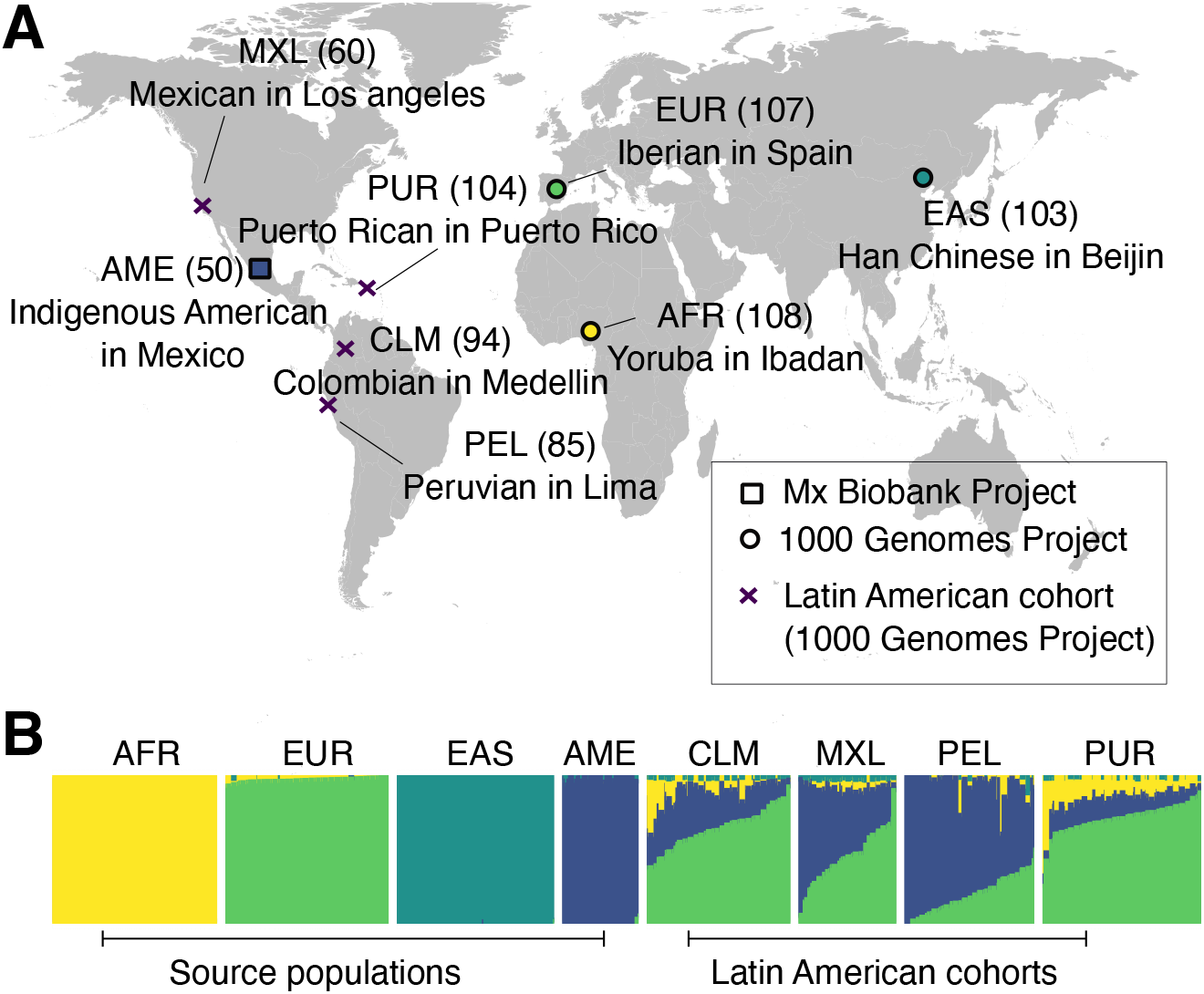
Populations used for demographic reconstruction. (A) Map showing the approximate sampling locations and the population names and codes. Numbers in parentheses denote the number of sampled genomes. Latin American populations are marked with a purple cross. (B) We used *ADMIXTURE* (with K=4) to visualize ancestry proportions of source populations and those from Latin America. The plot reveals substantial variation in ancestry proportions both across populations and among individuals within populations.

Our inferred four-population demographic model builds upon previously inferred models for the out-of-Africa expansion involving AFR, EUR, and EAS populations^7, 17, 23^. To include the Indigenous American population, we considered models in which their ancestors branched from those of EAS, introducing three additional parameters: the AME split time from EAS, the bottleneck size, and the population expansion rate (Figure 2A). We used *moments*^23^ to fit our parameterized models to the observed jSFS for three putative neutral mutation classes: intergenic, intronic, and synonymous variants, which were fit separately. Our inferred models for deep population history were consistent with the observed SFS for all three mutation classes (Figure 2B, C, and Figure S1, Table S1). The estimated 95% confidence intervals for the parameters overlapped across the three sets of SNPs (Figure 2C, Table S1). This consistency across annotations suggests that our inferred models are robust to differences in selective effects (either direct or linked) between mutation classes. We also evaluated alternative models of population size changes in the AME branch, but these models did not provide a better fit to the data (Figure S2). We found that the inferred AME split time was consistent across all tested models, with values ranging from 31,000–33,000 years ago.

**Figure 2.**
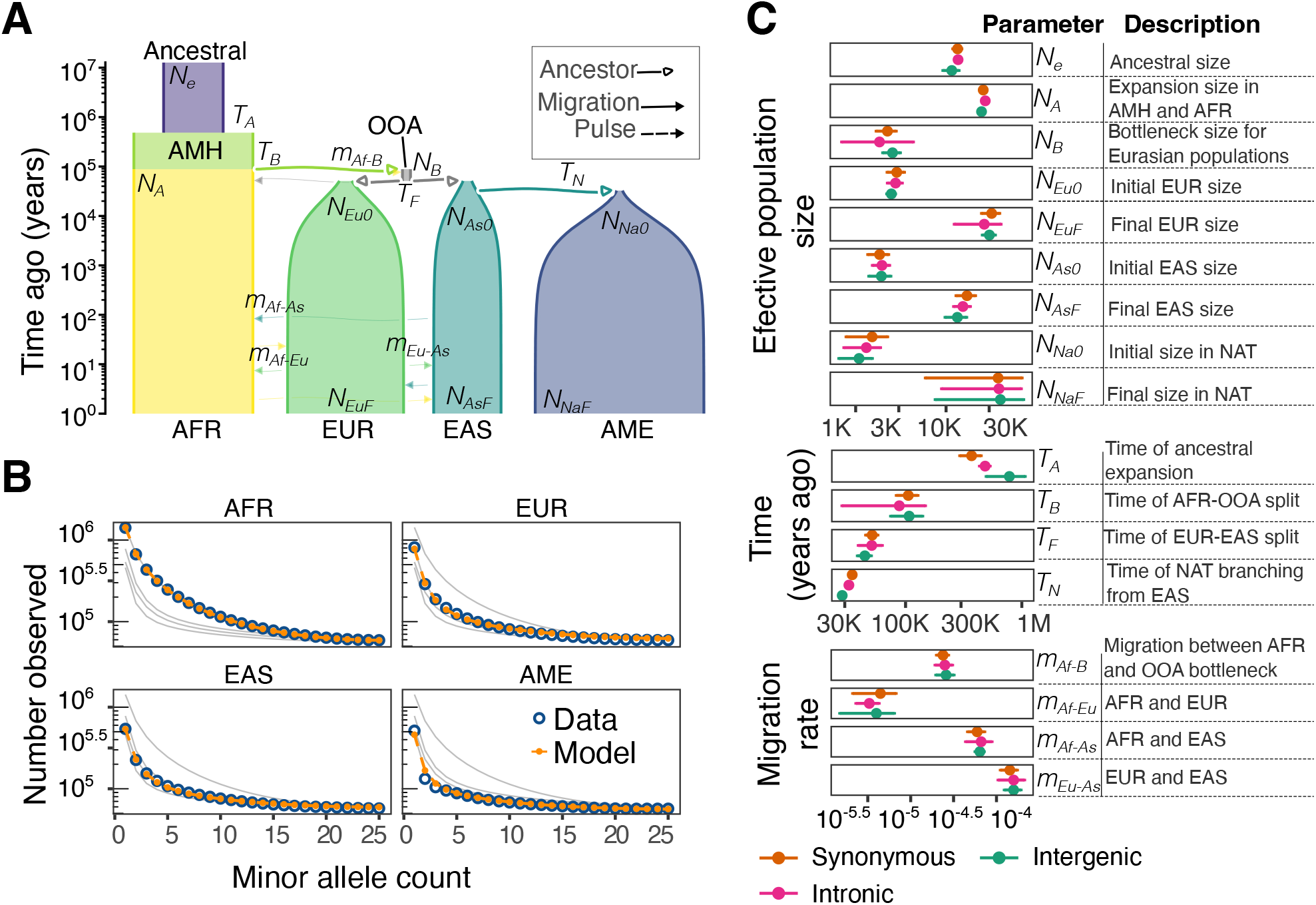
Inference of continental population history from allele frequencies. (A) Scheme of the inferred model. The width of the tubes is proportional to the effective population size; fonts in italics show the name of the inferred parameters. Abbreviations: Anatomically Modern Humans (AMH) and out-of-Africa (OOA). (B) Comparison between the folded intronic SFS for the data (blue) and the model’s prediction (orange). Background lines (grey) show the model’s prediction in the rest of the populations. (C) Comparison of estimated parameters with different sets of SNPs (synonymous, intergenic, and intronic). The error bar denotes the bootstrap confidence intervals (±2 standard deviations). The 95% confidence intervals overlap between annotations for the vast majority of inferred parameters.

While previously reported models have either excluded the Indigenous American population^8, 17^ or one or more of the other populations (European, African, and East Asian)^7, 12^ due to methodological limitations, we reconstructed a broad-scale demographic model for global human dispersal that includes the peopling of the Americas. By including all four populations in our analysis, we were able to provide a model that better predicts allele frequencies in each of these groups.

### Inferring admixture dynamics in Latin American populations

The recent history of Latin American populations involves multiple periods of immigration and extensive gene flow among different Indigenous American, European, and African populations^4, 16, 18^. Population genetic methods that rely on the jSFS may not be appropriate for resolving such admixture dynamics, as they typically assume a straightforward admixture process involving only a few ancestral populations and lack the necessary resolution to accurately infer the timing of admixture events that occurred relatively recently. However, by analyzing the ancestry tract length distribution (i.e., the distribution of lengths of continuous ancestry segments from each source population in admixed genomes), we inferred admixture models that include up to four ancestral populations and complex migration events^24^.

Using the distribution of ancestry tract lengths, we inferred admixture histories for four cohorts: Mexico (MXL), Puerto Rico (PUR), Colombia (CLM), and Peru (PEL). To calculate the ancestry tract length distribution, we constructed a reference panel of African, European, East Asian, and Indigenous American ancestries to perform local ancestry inference^25^ (Figure 3A and Methods). We treated each population independently, as admixture histories vary between regions in Latin America^4, 14, 16^, leading to diverse ancestry patterns (Figure 3B). In each population, we used *tracts*^24^ to test five admixture models including three source populations: AFR, EUR, and AME (Figure S3 and Methods). These models allowed us to explore a variety of admixture scenarios including single pulses of gene flow, multiple pulses, and continuous migration.

**Figure 3.**
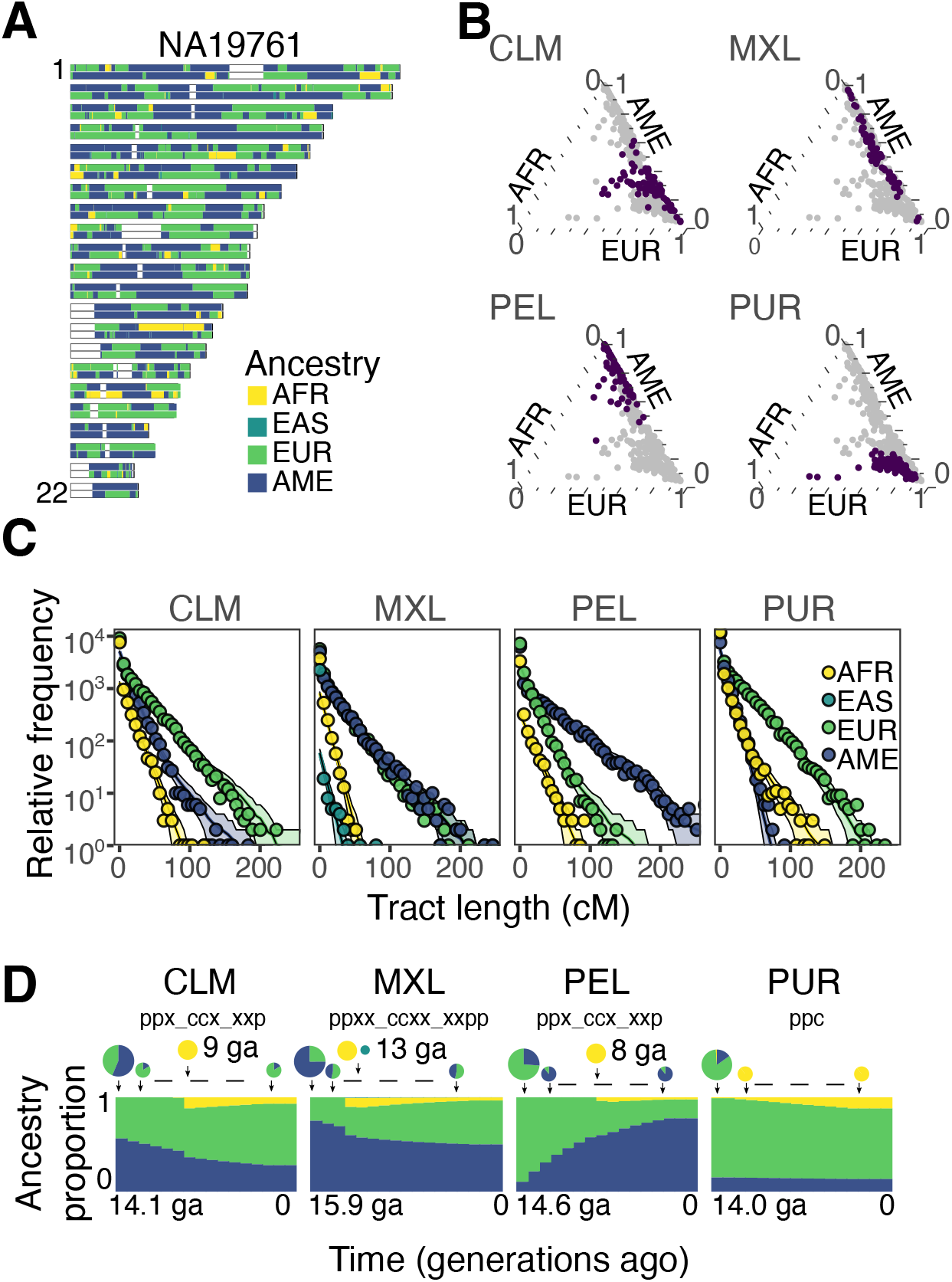
Inference of admixture history from the ancestry tract length distribution. (A) Karyogram with inferred local ancestry tracts in a Mexican individual (MXL). (B) Ternary plot of ancestry fractions for African, European, and Indigenous American ancestries inferred with *ADMIXTURE*. Points in purple correspond to the individuals from the population shown; grey points, on the background, are the other populations. (C) Ancestry tract length distribution of data and best model’s prediction. Plotted points show the aggregate tract length counts; lines show the maximum-likelihood k>est-fit tract length distributions for the best model in each population; and shading shows one standard deviation confidence interval, assuming a Poisson distribution of counts per bin. (D) Scheme of inferred admixture models. Top: pie charts sizes indicate an approximate proportion of migrants in each generation, and the pie parts represent the fraction of migrants of each origin in a given generation. Migrants are taken to have uniform continental ancestry. The dashed lines connecting the smaller circles denote a continuous migration similar to the connecting circles. Bottom: the y-axis shows the expected fraction of ancestry in the population at each point in time, and the x-axis represents generations ago, 15.9 ga corresponds to c: 1548, and 14 to c: 1604 (1 generation = 29 years).

We used a Bayesian Information Criterion approach (BIC) to compare the fit of these models and select the best-fit model (Figure 3C,D and Figure S4). For Colombia, the model selected based on the BIC did not correspond to known historical events, as the initial admixture event was too recent (~ 10 generations ago). In this case, we used additional information about known historical events to refine the model selection and chose a model that had a good fit to the data and parameters that were more consistent with known history^26, 27^. The models we selected (times of admixture events and ancestry proportions) are consistent with previous studies where admixture histories in Latin America have been explored^28–30^.

Previous work has identified genetic connections between Mexico and Asia that originated during the Manila galleon trade between the colonial Spanish Philippines and the Pacific port of Acapulco^15^. We identified ~ 1% (0.5 – 1) EAS ancestry among individuals in the MXL cohort using both *ADMIXTURE* and local ancestry inference. To incorporate a fourth source ancestry, we added an East Asian component to our admixture model for Mexico. We adapted our best-fitting three-population model to include an additional migration pulse from East Asia, occurring at the same time as the African pulse. This choice was motivated by the historical timing of the Manila galleon trade, which coincided with the period of the African slave trade^31, 32^. Our estimates indicate that this pulse occurred 13 generations ago (Figure 3C,D), which is consistent with published estimates^15^. We did not explore four-source population models for the other cohorts (CLM, PEL, and PUR), as we observed lower levels of East Asian ancestry in these populations using *ADMIXTURE* and local ancestry inference (Figure 1).

### Integrating Allele Frequencies and Ancestry Tracts: A Joint Modeling Approach for Demographic Reconstruction in Latin America

We combined our inferences from allele frequencies (Figure 2) and ancestry tracts (Figure 3) to provide a comprehensive model for the span of demographic history in Latin America (Figure 4A). Information from different inference engines (*tracts* and *moments*) was integrated using *demes*^33^, a standard format for demographic models in population genetics. In our combined model, we made several assumptions to simplify the analysis and produce a more tractable model. We assumed that the Latin American cohorts (MXL, CLM, PEL, and PUR) are independent of one another and have not experienced recent gene flow between them. We also fixed the effective population sizes (*N_e_*) in admixed populations to a constant value of 20,000 that roughly reflects the present-day *N_e_* in Latin American populations^8^. We did not infer *N_e_* in these populations because the ancestry tract distribution does not contain information about *N_e_*, and the admixture histories in Latin America are very recent on an evolutionary time scale so that allele frequencies will not fluctuate significantly due to genetic drift.

**Figure 4.**
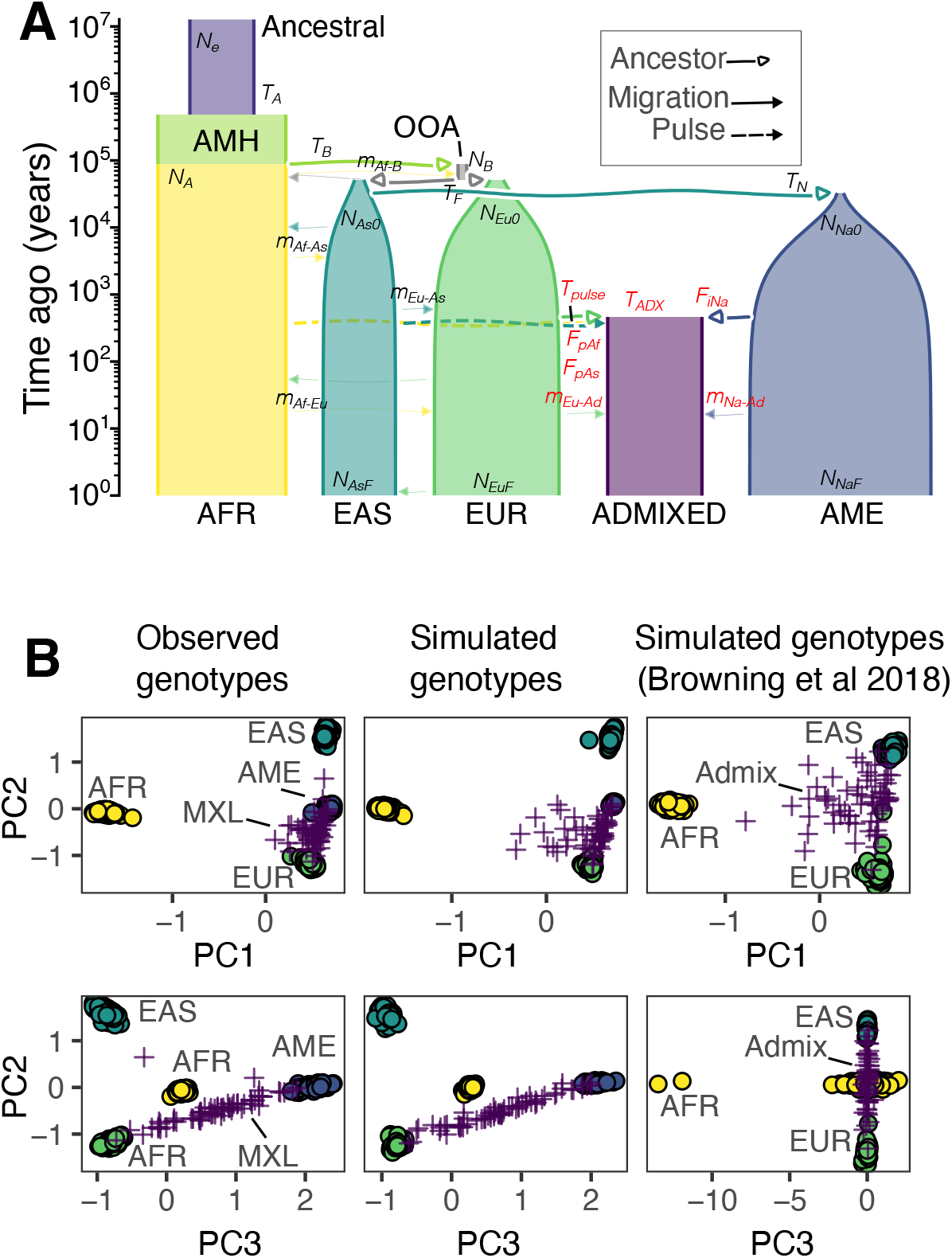
Model incorporating both time scales of population history, continental history and admixture dynamics. (A) Visualization of the inferred demographic model. We combined the inferences from allele frequencies and ancestry tracts into a single demographic model. The parameters in red correspond to the parameters estimated from ancestry tracts and the parameters in black were inferred from allele frequencies. The admixed population, in purple, represents the Mexican population (MXL). (B) PCA decomposition of genetic data. Left: data chromosome 22; middle: simulation using our inferred model (panel A); right: simulation using the Browning *et al*. model. For both models, we simulated a genome of length 50Mb with *msprime* (Methods). In the data (bottom left) the Indigenous American similarity component is capture by the PC3. This variation is capture in our inferred model (A), but is not present in the simulated genomes from the Browning *et al*. model, which uses the East Asian population as a proxy for Indigenous American ancestry.

To validate the accuracy of our joint demographic model, we used simulations to generate artificial datasets and compare them to the observed data. We used the coalescent simulator *msprime*^34, 35^ to simulate samples for each population under the reconstructed demography (Methods). We compared a Principal Component Analysis (PCA)between the simulated data and observed genotypes on chromosome 22 (Figure 4B). The PCA of simulated individuals closely mirrors those in the observed data (Figure S5), indicating that our model accurately captures broad-scale patterns of genetic diversity in Latin American populations.

To compare to existing models for admixed Latin American populations, we also simulated data under the Browning *et al*. (2018) model of American admixture^6, 8^. Using the same simulation and PCA approach, we compared a PCA between this model and real data, finding that our model (Figure 4B) more accurately recapitulates important axes of genetic diversity in Latin American populations. In particular, our model reproduces the variation in Indigenous American ancestries among individuals, as shown by the strong association between the principal component 3 (PC3) and Indigenous American ancestries. In contrast, the Browning *et al*. model has no association with Indigenous ancestries in PC3, as it uses East Asian ancestry as a proxy for American ancestry.

### Applying our demographic model to the simulation of functional variation under selection

To evaluate the ability of our inferred demographic model to explain patterns of genetic variation under selection, we conducted forward-in-time simulations of non-coding (intergenic and intronic), synonymous, missense, and loss-of-function variants using the *fwdpy11* simulation engine^36, 37^. We assessed the ability of our demographic model, when combined with inferred models of selection, to explain patterns of genetic variation in different functional categories of the genome.

Missense and loss-of-function (nonsense) mutations are often subject to negative selection because they can have significant impacts on protein function. While non-coding and synonymous variants may be under direct selection due to changes in regulatory elements and codon usage bias^38^, respectively, here we assumed that they were neutral with respect to selection^39, 40^. To simulate selection on nonsynonymous mutations, we incorporated distributions of fitness effects (DFE) for the two classes of nonsynonymous mutations^41^ (Figure S6A and Methods). We simulated a total of 350Mb of genomic data using our inferred demographic model and spanning various regions of the human genome by sampling from existing recombination maps and genomic annotations (Methods). Due to the computational burden of running forward-in-time simulations, we limited the simulation to only one Latin American population (MXL) jointly with the other source populations (AFR, EUR, EAS, and AME).

We evaluated how well our joint modeling approach matched data from selected mutation classes and assessed any biases within putatively neutral classes due to linked selection. We found that the SFS from simulated data accurately reproduced the SFS from the data for both neutral and selected genetic variation (Figure 5A and Figure S7). Specifically, the SFS fits well for non-coding, synonymous, and missense variants, for which many variants are observed. While there are fewer observed loss-of-function variants, leading to noisier estimates of the SFS, simulated data is largely unbiased across frequency bins (Figure S7). In addition to analyzing the SFS within single populations, we calculated pairwise *F*_ST_ between populations. We observed that the simulated *F*_ST_ values accurately reproduced those from the observed data (Figure 5B). Together, this modeling approach was able to capture observed patterns of allele frequency variation within and between populations, supporting the utility of our model as a tool for studying complex evolutionary processes in these cohorts through genetic simulations.

**Figure 5.**
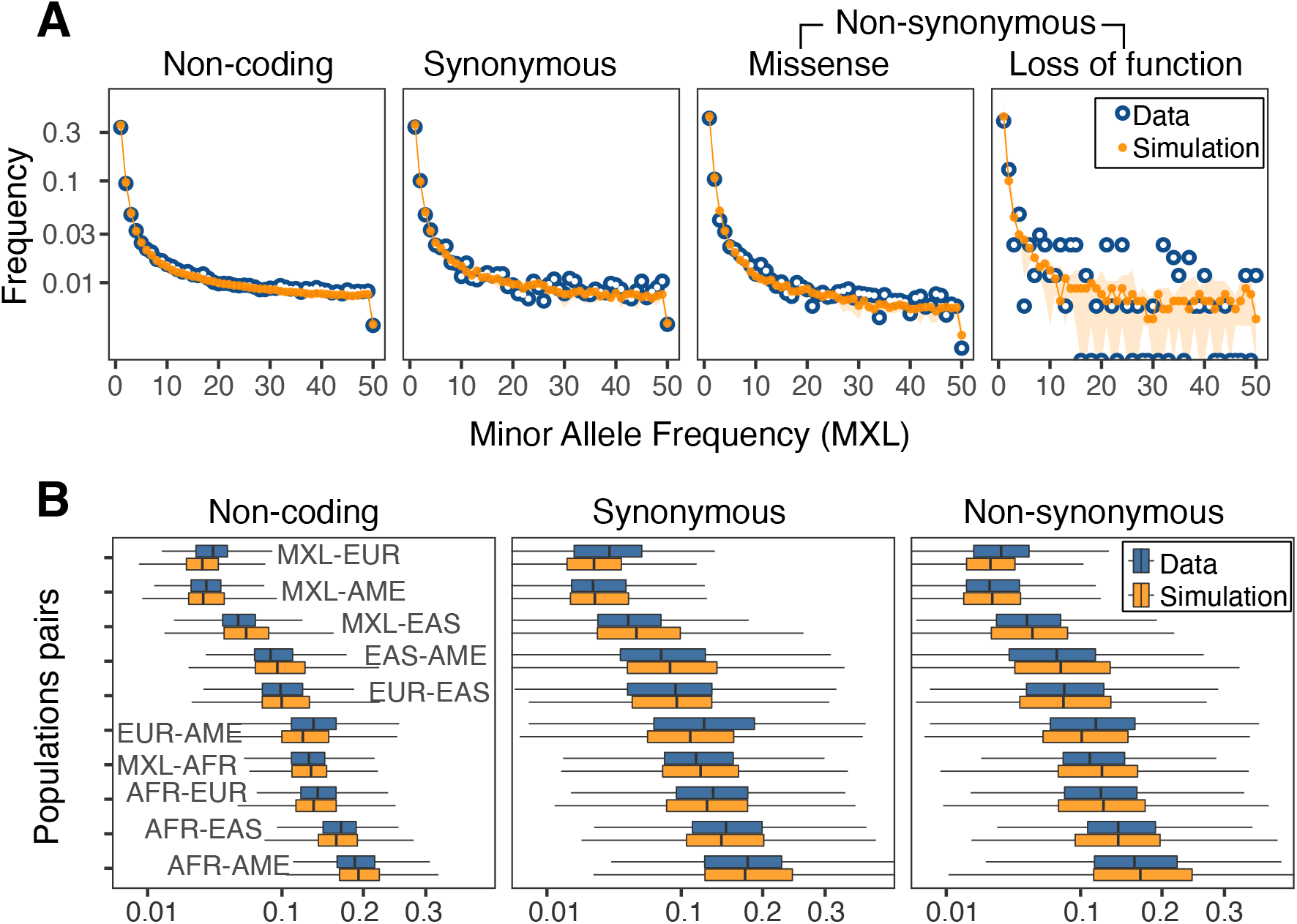
Forward-in-time simulations of functional genetic variation. We simulated a total of 350 independent regions, 1Mb each, of the human genome by sampling coding annotations and inferred recombination rates (Methods). (A) Comparison between the folded SFS for the data (blue) and the simulation (orange) in different functional categories, summed across all regions. The SFS corresponds to the admixed Mexican population (MXL). Additional populations are shown in Figure S7A. (B) Box-plot of pairwise *F_ST_* values comparing data (n = 350) and simulation (n = 350). Missense and loss-of-function mutations were combined into a single non-synonymous category.

## Discussion

Our study provides comprehensive demographic models for populations in Latin America inferred from high-coverage whole genome data. These models allow us to examine population dynamics at different time scales, incorporating multiple epochs of demographic history. By inferring a four-populations Out-of-Africa model for the deeper demographic history (Figure 2), and using ancestry tract length distributions to infer the recent admixture history in four Latin American cohorts (Figure 3), we were able to create a single model (Figure 4) that jointly captures both allele frequencies and the distribution of ancestry tract lengths. Through extensive simulations, we demonstrated that this model accurately reflects the patterns of genetic diversity present in Latin American populations (Figure 5).

Our inferred demographic models for populations in Latin America represents a significant advance in our ability to accurately account for their evolutionary history in genomic studies. By capturing different aspects of population dynamics, our model provides a powerful tool for studying Latin American populations and understanding the genetic basis of disease in these underrepresented groups^42^. Populations and individuals with recent genetic admixture are routinely excluded from genomic studies due to concerns over population structure^43^. This is due in part to the lack of methods and pipelines to effectively account for shared ancestries and complex demographic histories^44–46^, which our study takes steps to address. Past demographic history can also pose a significant challenge to detecting selection, as it can create patterns of genetic diversity that mimic the signatures of selection^47–49^. Detailed demographic models are important for controlling for such confounding, allowing for more robust inference of recent selection in Latin American populations. Our models therefore provide a resource for developing and testing methods and pipelines that are specifically tailored to studying admixed populations, improving the accuracy and validity of results obtained from such studies.

Our results reveal a dynamic and multifaceted history. Of note, the inferred AME-EAS split time ranges from 31,000 to 33,000 years ago and is consistent across all tested models. This suggests that population structure between the ancestors of Indigenous Americans and East Asians had formed by this time. This may support the idea of an early migration to the Americas, predating the commonly used time frame of ~ 15,000 years ago, or reflect population structure in Siberia and Beringia prior to the expansion into the Americas^50, 51^. Additionally, our results show that the ancestry proportions in Latin American populations have changed over time, with evidence of continuous migration of European and Indigenous ancestries in Mexico, Peru, and Colombia. Similar patterns of changes in ancestry proportions over time have been observed in other admixed populations in the United States^52^. This suggests that there may have been ongoing migrations from Europe to the Americas, as well as migration within the Americas, such as the movement of people from rural to urban areas^53, 54^. This highlights the importance of considering the heterogeneity of population histories and the historical and cultural interactions of the Americas in understanding the demographic history of Latin American populations.

Our models provide valuable insights into the larger-scale demographic history of Latin American populations, but they do not capture all aspects of population history. For example, the arrival of the Spanish in central Mexico led to a significant decline in population size^55^, which can have a significant impact on genetic diversity but is not fully captured by our model. With the increasing availability of larger datasets, it will be possible to study more detailed population structure and dynamics using the framework from this study. As we continue to increase the representation of Latin American genomic data, including more whole genomes, we can expect to gain a deeper understanding of the complexity of Latin American population history and to resolve historical processes at finer scales.

## Methods

### Whole genome data and quality control

We used two high-coverage whole genome datasets to investigate the demographic dynamics of Latin American populations. The first dataset was the 1000 Genomes Project (1KGP)^22^, which provided 30x high-coverage genomic data from the NYGC that was mapped to the GRCh38 reference genome. We selected reference samples for Africa (AFR), Europe (EUR), and East Asia (EAS) from this data set, specifically we used 108 Yorubans in Ibadan Nigeria, 107 Iberians in Spain, and 103 Han Chinese in Beijing, respectively. In addition to these reference samples, we also used data from admixed cohorts within the 1KGP, including 60 Mexicans in Los Angeles (MXL), 104 Puerto Ricans in Puerto Rico (PUR), 94 Colombians in Medellin (CLM), and 85 Peruvians in Lima (PEL), to infer admixture history. The second data set was from the MX Biobank (MXB) project^3,20^ and included 50 high-coverage whole genomes from Indigenous American (AME) descendants in present day Mexico.

To combine the data sets from the 1KGP MXB projects, we used the *CrossMap*^56^ tool to liftover the MXB genomes from the GRCh37 reference genome to the GRCh38 reference genome. Next, we applied two rounds of filters to remove variants that were likely affected by genotyping error or batch effects. The first filter, applied to each source population (AFR, EUR, EAS, and AME) separately, removed variants in Hardy-Weinberg diseqeuilibrium with a p-value of 1e-4. The second filter, with a more strict Hardy-Weinberg p-value of 0.05, was applied with the purpose of removing potential batch effects. In this case, we applied the second filter to the 50 MXB genomes combined with a set of 29 individuals from the 1KGP (2 MXL and 27 PEL) with the highest proportion of Indigenous American ancestries. In addition to these filters, we also removed variants that were located in masked regions of the genome. After applying these filters, we obtained a combined data set that was used for all downstream analyses in the study.

### Admixture analysis

We used the software package *ADMIXTURE* v1.3^57^ to estimate individual ancestry proportions based on genome-wide single nucleotide polymorphism (SNP) data. We applied the following filters to the data: SNPs with a minor allele frequency below 0.05 and SNPs with a Hardy-Weinberg equilibrium test p-value below 0.001 were excluded. In addition, we filtered SNPs in linkage disequilibrium with each other using a threshold of *r*^2^ = 0.1. We ran *ADMIXTURE* with default parameters, specifying the number of ancestral populations (K) to range from 2 to 6.

### Demographic inference from allele frequencies

To accurately infer the demographic model, we excluded variants located in CpG sites, which have been shown to have highly variable mutation rates and could potentially bias the results^58, 59^. To infer the demographic model, we calculated the folded joint site frequency spectrum (jSFS) for intronic, intergenic, and synonymous variants. We classified the variants according to its functional effect with *VEP* version 102^60^. We down sampled to sample sizes of 40 individuals from each population, resulting in a jSFS with four dimensions: one for each of the four populations (AFR, EUR, EAS, and AME).

To infer demographic parameters, we used *moments* as our inference engine due to its ability to handle inferences for more than three populations using a diffusion approximation^23^. We adopted a two-step approach to inferring the demographic parameters. In the first step, we inferred a 3-populations out-of-Africa model (AFR, EUR, and EAS)^7, 23^, which included 14 parameters such as effective population sizes, split times, and migration rates. In the second step, we included the Indigenous American population as branching out from the ancestors of the East Asian population and inferred three additional parameters: the AME split time from EAS, the bottleneck size, and the population expansion rate (Figure 2A). We performed this same inference for each class of putatively neutral jSFSs (intronic, intergenic, and synonymous). To account for uncertainty in the parameters, we divided the genome into non-overlapping windows of 10Mb and calculated the jSFS and scaled mutation rates for each window. We generated bootstrap replicates from these windows (*n* = 288) by sampling with replacement and used a Godambe Information approach^61^ to estimate parameter uncertainties.

### Estimation of mutation rates

To convert genetic to physical units (i.e., event times in generations, and effective instead of relative population sizes) we required estimates of total mutation rates in retained genomic regions. To estimate this scaled mutation rate, *μL*, for non-coding (intronic and intergenic) and coding (missense, synonymous, and loss of function) regions of the genome, we first subset the genomic coordinates for these regions. For intergenic and intronic regions, we counted the occurrence of each triplet context (excluding CpG contexts) and used the corresponding total mutation rates for each possible mutation for each triplet context, obtained from the gnomAD mutation model^58^, to weight the counts. We then summed the weighted counts for all triplets to estimate the scaled mutation rate for non-coding regions. For coding regions, we generated a file with every possible mutation (biallelic SNPs) in the coding regions of the genome and used *VEP* version 102 tool to predict the consequence of each variant^60^. For each consequence (synonymous, missense, or nonsense), we counted the occurrence of each triplet and used the same approach as for non-coding regions to estimate the scaled mutation rate.

### Local ancestry inference

To infer the ancestry of specific ancestry segments within the Latin American genomes, we utilized *Gnomix*^25^. In order to do so, we first constructed a reference panel consisting of individuals with known African, European, and Indigenous American ancestries. Particularly, we selected the Yoruba population (YRI=108) to represent African ancestries, the Iberian population in Spain (IBS=107) and the British in England and Scotland (GBR=91) for European ancestries, and a combination of MXB Indigenous American samples (n=50) with 30 1KGP individuals with high Indigenous American ancestries proportions from Peru (PEL=28) and Mexico (MXL=2) for Indigenous American ancestries. In addition, we also generated a reference panel including East-Asian ancestry, using the Han Chinese population (CHB=103) as a reference. We then trained two *Gnomix* models using these two reference panels, using the default settings which are optimal for whole genome data and setting the phase parameter to True, in order to re-phase the genomes using the predicted local ancestry. Finally, we used these models to predict the local ancestry in each of the Latin American genomes.

### Inference of admixture history

To infer the admixture history, we used the ancestry tract length distribution obtained from the *Gnomix* output, which we collapsed into haploid ancestry tracts. We analyzed the admixture dynamics independently in each Latin American cohort (MXL, PEL, CLM, and PUR). We evaluated five different admixture models with *tracts*^24^: *ppx_xxp, ppx_xxp_pxx, ccx_xxp, ppx_ccx_xxp*, and *ppc*. The order of each letter in these models corresponds to the order of the source ancestries (AMR, EUR, and AFR), with an underscore indicating a distinct migration event. A “*p*” represents an ancestry pulse (e.g. *xxp* is a pulse from AFR ancestry), “*c*” indicates continuous migration, and “*x*” indicates no input. We selected these models for analysis because they cover a range of plausible admixture history scenarios and have been previously tested in similar cohorts^28–30, 62^.

To account for the uncertainty in estimated parameters, we generated 100 bootstrap replicates of the ancestry tract distribution by randomly sampling individuals with replacement. For each replicate, we calculated the Bayesian Information Criterion (BIC) to determine the best-fit model. The selected model for each population was based on the BIC as well as agreement with historical evidence. In the case of Mexico, we used the *ppx_ccx_xxp* model as the best fit and added a pulse of Asian ancestry occurring simultaneously with the African pulse. We optimized this model by exploring the parameter space to find the best fit parameters (modeled as *ppxx_ccxx_xxpp*).

### Combining tracts and moments inferences

We integrated our demographic inferences from allele frequencies and tracts into a unified model. To do so, we utilized *demes*33 to combine the parameters. In order to account for variations in the generation time, we used a value of 29 years^63^. This enabled us to transform the parameters of the tracts (e.g. the timing of admixture events) from generations to years from present.

### Neutral simulations

To ensure the accuracy of our demographic model, we conducted coalescent simulations of neutral genetic sequence data. We used *msprime*^34, 35^ to implement the classical coalescent with recombination model (Hudsons algorithm) for the majority of the simulation, and for the last 20 generations we employed the discrete-time Wright-Fisher model^64^, which better accounts for large sample sizes and genome lengths and provides more accurate patterns of identity-by-descent and ancestry tract variation in admixed populations. To further verify the correctness of our model, we utilized multiple simulation engines to compare to observed data, including *msprime, fwdpy11*, and *moments*. We utilized *tskit* for downstream analysis of the simulated genomes.

### PCA analysis

We performed a principal component analysis (PCA) on three datasets using the *scikit-allel* software^65^. The three datasets were: (1) observed genotypes, (2) simulated genotypes generated by our model, and (3) simulated genotypes generated by the Browning *et al*. model^8^. For the PCA analysis, we excluded all singleton variants. We also performed linkage disequilibrium (LD) pruning by removing variants in high LD (*r*^2^ > 0.1). The PCA analysis was conducted separately for each dataset.

### Forward-in-time simulations

We performed forward simulations using the software *fwdpy11* version 0.18.3^36, 37^. Our simulations incorporated our inferred demographic model. To ensure the realism of our simulations, we divided the human genome (autosomes only) into windows of 1Mb and randomly selected 350 of these regions for simulation. We then analyzed the functional annotation of each of these regions, including intergenic, intronic, and coding sequences, and calculated the corresponding mutation rates. To accurately capture the effects of recombination on evolutionary dynamics, we incorporated a human recombination map for GRCh38 obtained from https://github.com/odelaneau/shapeit4/tree/master/maps into our simulations^66^. We incorporated distributions of fitness effects (DFE) for missense and nonsense mutations, in which nonsense mutations were inferred to be more strongly selected against^41^. To do this, we employed a multiplicative model that allowed us to examine the influence of selective pressures on the genetic diversity of our simulated populations (Figure S4). In comparing between simulations and data, we aggregated data across the 1Mb regions of the genome that were simulated. The simulation was run for 152,471 generations, corresponding to the recent history of the five modeled populations. Finally, we sampled individuals from each of the five populations at the present time, with sample sizes matching those of our data.

## Data availability

The 1KGP data^21, 22^ is publicly available and accessible without restriction. The 50 genomes from the MX-Biobank^3, 20^ project are deposited in the European Genome-phenoms Archive (EGA) repository, accession number EGAD00001008354.

## Demographic models

The inferred demographic models are provided as supplementary material and in the GitHub repo: https://github.com/santiago1234/mxb-genomes.

The model files are in the *demes*33 format.

- **Model 1** (*m1-out-of-africa.yml*). Four populations out of Africa.
- **Model 2** (*m2-Mexico-admixture.yml*). Admixture in Mexico.
- **Model 3** (*m3-Colombia-admixture.yml*). Admixture in Colombia.
- **Model 4** (*m4-Peru-admixture.yml*) Admixture in Peru.
- **Model 5** (*m5-PuertoRico-admixture.yml*). Admixture in Puerto Rico.
- **Model 6** (*m6-All-admixture.yml*). Admixture in Latin American populations, combined across models 2–6.

## Code availability

Code for the software used in this paper is found at the following locations: *moments* (https://bitbucket.org/simongravel/moments), *tracts* (https://github.com/sgravel/tracts), *demes* (https://github.com/popsim-consortium/demes-python), *msprime* (https://github.com/tskit-dev/msprime), *tskit* (https://github.com/tskit-dev/tskit), *fwdpy11* (https://molpopgen.github.io/fwdpy11/), and code used to process data, infer demographic models, run simulations, and generate visualizations is available at https://github.com/santiago1234/mxb-genomes.

## Acknowledgements

This work was supported by “The Mexican Biobank Project: Building Capacity for Big Data Science in Medical Genomics in Admixed Populations”, a bi-national initiative between Mexico and the UK co-funded equally by CONACYT (Grant number FONCICYT/50/2016), and The Newton Fund through The Medical Research Council (Grant number MR/N028937/1) awarded to A.M.-E. Both A.P.R. and S.G.M.-M. were financially supported by CONACYT with funds from the MX Biobank Project and a graduate program scholarship, respectively. We also thank Carmina Barberena Jonas and Ram Gonzales for their feedback throughout the project and Jacob Cervantes for IT support. We also would like to acknowledge Kevin Thornton for his assistance with setting up the forward-in-time simulations using *fwdpy11*.

## Author contributions statement

A.P.R. and A.M.-E. conceived the study and provided overall supervision. S.G.M.-M. and A.P.R. carried out the analyses and were responsible for writing the initial manuscript. D.O.V. and A.M.-E. provided feedback on the manuscript. L.G.G., L.P.C.-H., and L.F.-R. contributed to the acquisition of the data. All authors read and approved the final manuscript.

## Additional information

### Competing interests

The authors declare no competing interests.

## Notes

https://github.com/santiago1234/mxb-genomes

### Competing Interest Statement

The authors have declared no competing interest.

