## Supplemental figures and tables for "Demographic Modeling of Admixed Latin American Populations from Whole Genomes"

SUPPLEMENTAL TABLES AND FIGURES FOR:
DEMOGRAPHIC MODELING OF ADMIXED LATIN AMERICAN POPULATIONS FROM WHOLE GENOMES

| Description |  | Intergenic |  | Intronic |  | Synonymous |  |
| --- | --- | --- | --- | --- | --- | --- | --- |
|  |  | Value | SE | Value | SE | Value | SE |
| Time event (Thousands of years ago) |  |  |  |  |  |  |  |
| $T_A$ | Ancestral expansion | 782.167 | 144.204 | 483.457 | 25.421 | 369.958 | 38.717 |
| $T_B$ | AFR-OOA split | 107.416 | 16.526 | 88.348 | 29.993 | 105.570 | 11.170 |
| $T_F$ | EUR-EAS split | 44.669 | 3.075 | 51.166 | 5.976 | 51.622 | 3.120 |
| $T_N$ | EAS-AME branching | 28.516 | 0.517 | 32.555 | 0.514 | 34.758 | 1.002 |
| Effective population size (# of individuals) |  |  |  |  |  |  |  |
| $N_e$ | Ancestral size | 11,577 | 1,174 | 13,580 | 306 | 13,411 | 767 |
| $N_A$ | AMH and AFR size | 24,691 | 747 | 27,142 | 588 | 25,782 | 1,151 |
| $N_B$ | Bottleneck size | 2,557 | 283 | 1,835 | 1,260 | 2,247 | 280 |
| $N_{Eu0}$ | Initial EUR size | 2,483 | 138 | 2,761 | 259 | 2,847 | 309 |
| $N_{EuF}$ | Final EUR size | 30,163 | 2,480 | 26,462 | 7,067 | 31,955 | 3,648 |
| $N_{As0}$ | Initial EUR size | 1,924 | 263 | 1,955 | 211 | 1,840 | 240 |
| $N_{AsF}$ | Final EAS size | 13,339 | 1,792 | 15,364 | 1,617 | 17,038 | 2,073 |
| $N_{Na0}$ | Initial size in AME | 1,091 | 220 | 1,313 | 286 | 1,522 | 369 |
| $N_{NaF}$ | Final size in AME | 39,920 | 16,167 | 38,462 | 14,769 | 37,643 | 15,872 |
| Migration rate (Fraction of individuals per generation moving between populations) |  |  |  |  |  |  |  |
| $m_{Af-B}$ | AFR and OOA | $2.58 \times 10^{-05}$ | $3.11 \times 10^{-06}$ | $2.5 \times 10^{-05}$ | $2.89 \times 10^{-06}$ | $2.38 \times 10^{-05}$ | $1.92 \times 10^{-06}$ |
| $m_{Af-E}$ | AFR and EUR | $3.99 \times 10^{-06}$ | $1.25 \times 10^{-06}$ | $3.29 \times 10^{-06}$ | $4.98 \times 10^{-07}$ | $4.44 \times 10^{-06}$ | $1.17 \times 10^{-06}$ |
| $m_{Af-As}$ | AFR and EAS | $6.41 \times 10^{-05}$ | $3.93 \times 10^{-06}$ | $6.62 \times 10^{-05}$ | $1.13 \times 10^{-05}$ | $5.95 \times 10^{-05}$ | $6.74 \times 10^{-06}$ |
| $m_{Eu-As}$ | EUR and EAS | $1.59 \times 10^{-04}$ | $1.76 \times 10^{-05}$ | $1.59 \times 10^{-04}$ | $2.71 \times 10^{-05}$ | $1.43 \times 10^{-04}$ | $1.62 \times 10^{-05}$ |

**Table S1.** This table displays the inferred parameter values and the corresponding standard errors (SE) calculated from allele frequencies across different SNP categories. See also main figure 2A.

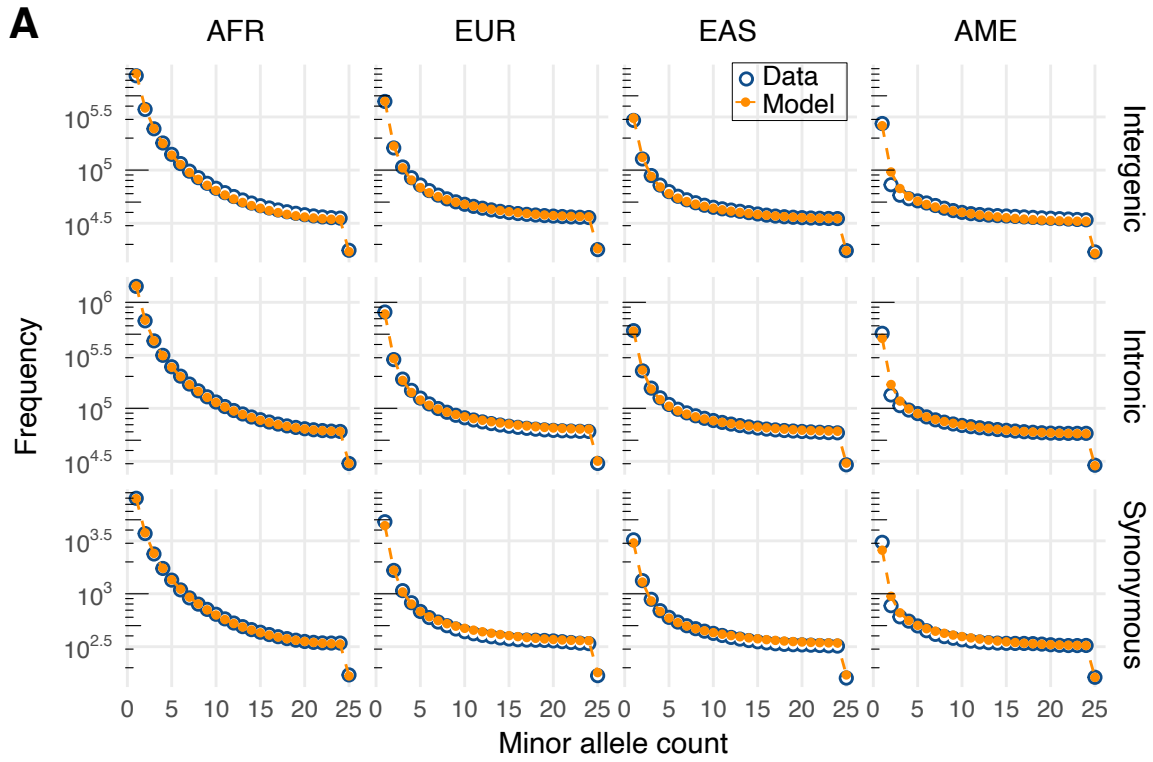

**Figure S1.** Comparing the Fits of Inferred Demographic Models across Different SNP Categories. Our analysis independently inferred demographic parameters for three sets of putative neutral variants: intergenic (top), intronic (middle), and synonymous (bottom). This figure presents a visual comparison between the observed folded site frequency spectrum of the data, depicted in blue, and the predicted site frequency spectrum generated by the demographic models, depicted in orange, across different populations.

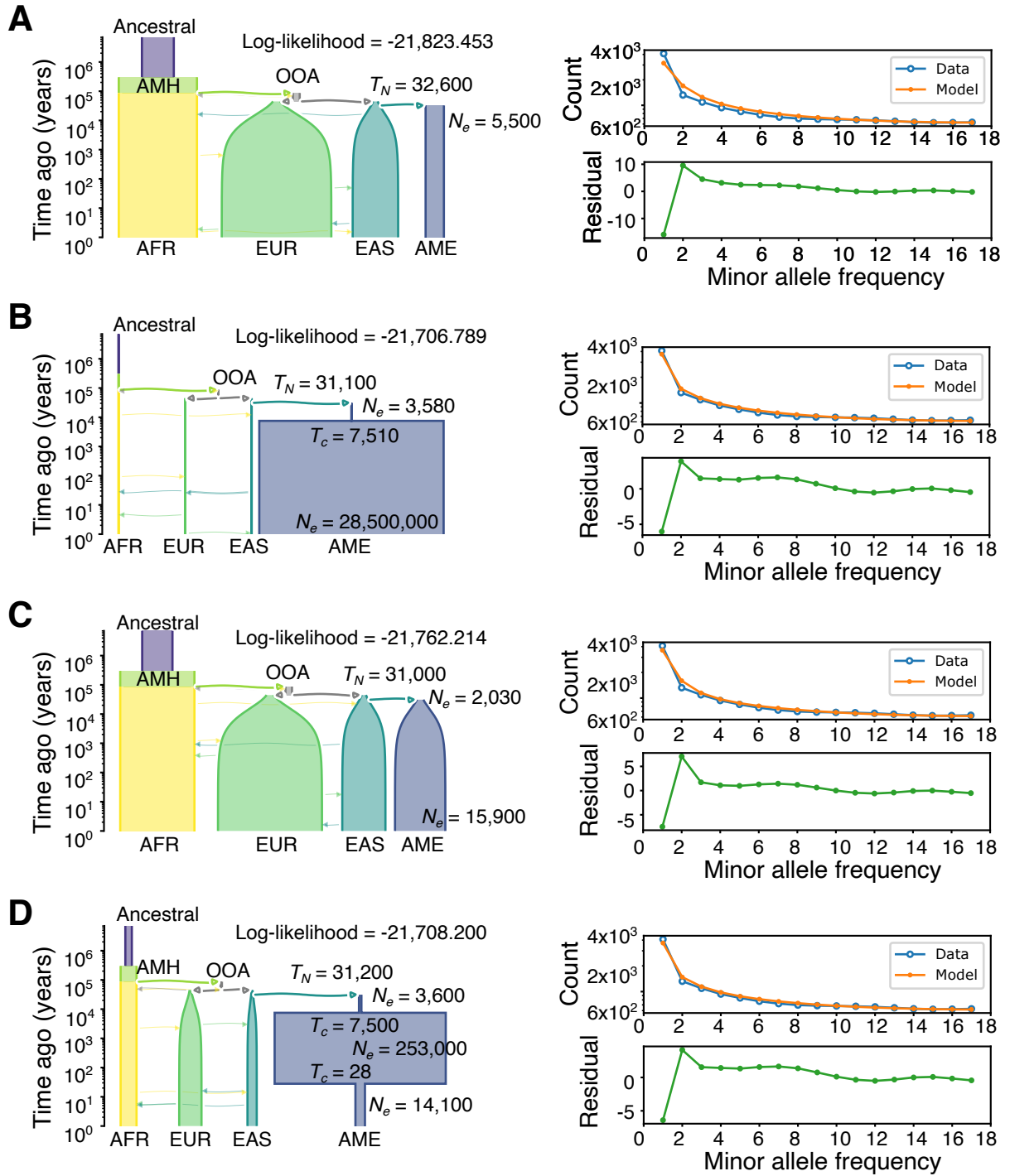

**Figure S2.** In this figure, we present a comparison of different demographic models for the Indigenous American population. From top to bottom, we consider a model of constant population size (A), a model of two epochs with constant population size in each (B), a model of exponential growth (C), and a model of three epochs with constant population size in each (D). The log-likelihood of each model is displayed in the top right corner, allowing us to assess the relative fit of each model to the data. The scheme of the inferred model is shown in the left, with the width of the tubes representing the effective population size and the italicized fonts indicating the name and values of the inferred parameters. The abbreviations AMH and OOA refer to Anatomically Modern Humans and Out-of-Africa, respectively. The plots on the right show a comparison between the folded intronic site frequency spectrum (SFS) for the data (shown in blue) and the model's prediction (shown in orange).

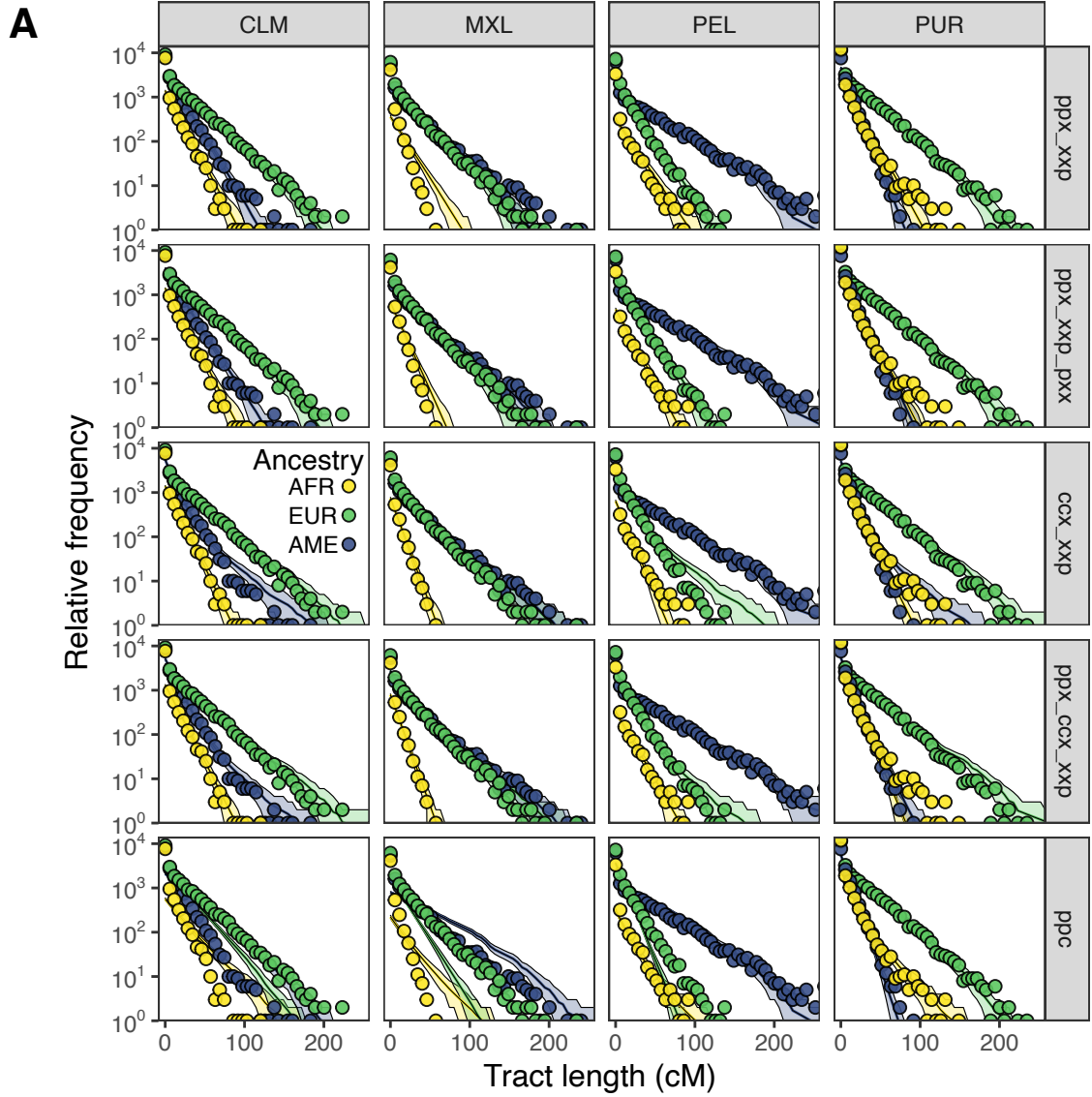

**Figure S3.** (A) In this figure, we compare the observed ancestry tract length distribution in Latin American populations to the predictions of the tested admixture models. The plotted points represent the data, while the lines show the model's predictions. The figure is organized by population (columns) and model (rows). The shading represents the uncertainty in the predicted distribution, with one standard deviation shown as a confidence interval assuming a Poisson distribution of counts per bin.

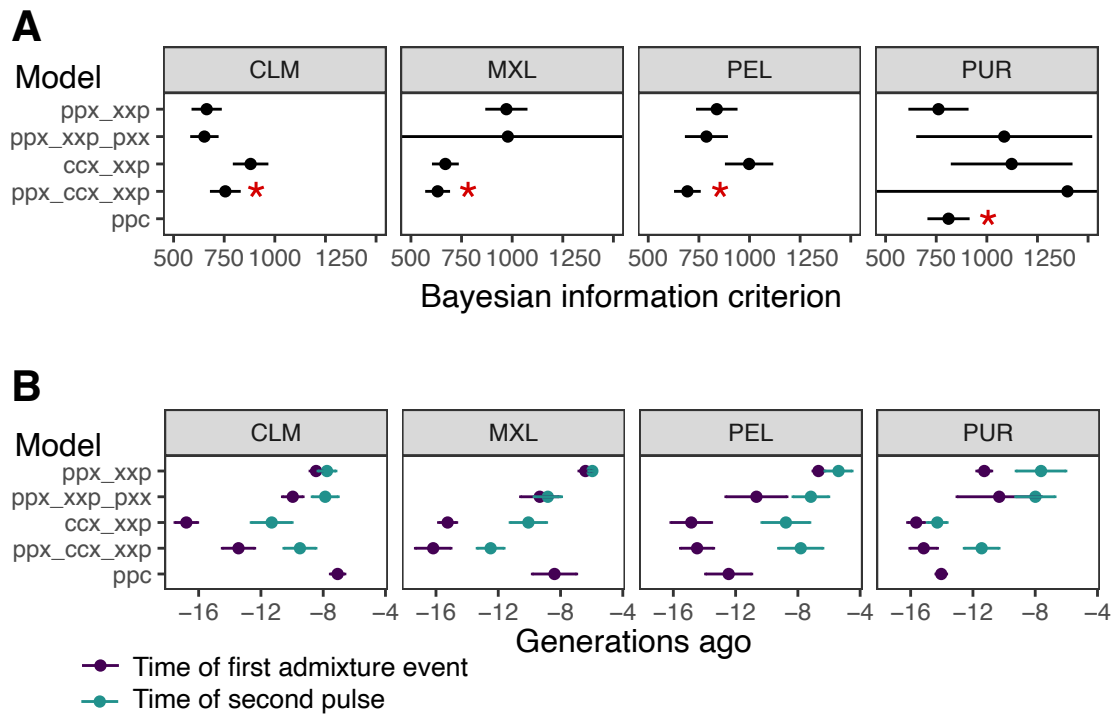

**Figure S4.** (A) The Bayesian Information Criterion (BIC) was calculated for each model, with uncertainty estimated through the generation of 100 bootstrap replicates of the data. The error bars shown represent $\pm 2$ standard deviations of the BIC values calculated from the bootstraps. The red star shows the model we selected. (B) All models have two common parameters: the time of the initial admixture event and the time of the African pulse (second pulse). These parameters are compared across the models in the figure.

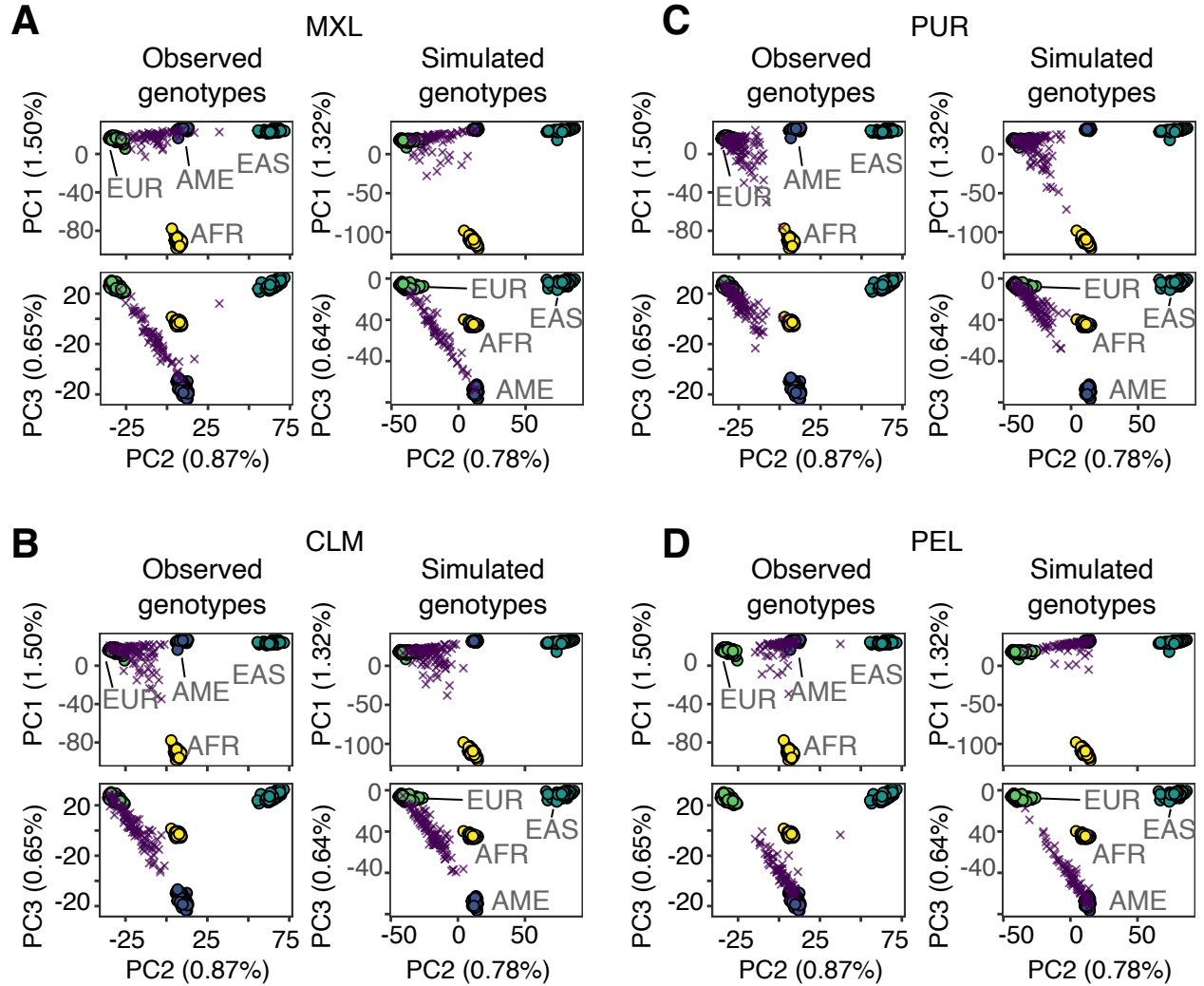

**Figure S5.** (A) We conducted simulations using our inferred demographic model and compared the results to observed genotype data. The PCA plots demonstrate a strong match between the observed and simulated data, with the observed genotypes plotted in the left column and the simulated genotypes plotted in the right column. The PCA analysis shows the distribution of ancestries in the data, with PC1 capturing the African ancestries component, PC2 capturing the European and East-Asian ancestries, and PC3 capturing the Indigenous American ancestries. The populations represented in the plot include those of African, European, East-Asian, and Indigenous American ancestries, shown in circles, and cosmopolitan individuals from Mexico, shown in purple crosses. (B) - (D) Same as (A) but showing the rest of the Latin American populations, purple crosses, the other populations, in circles, are always kept the same.

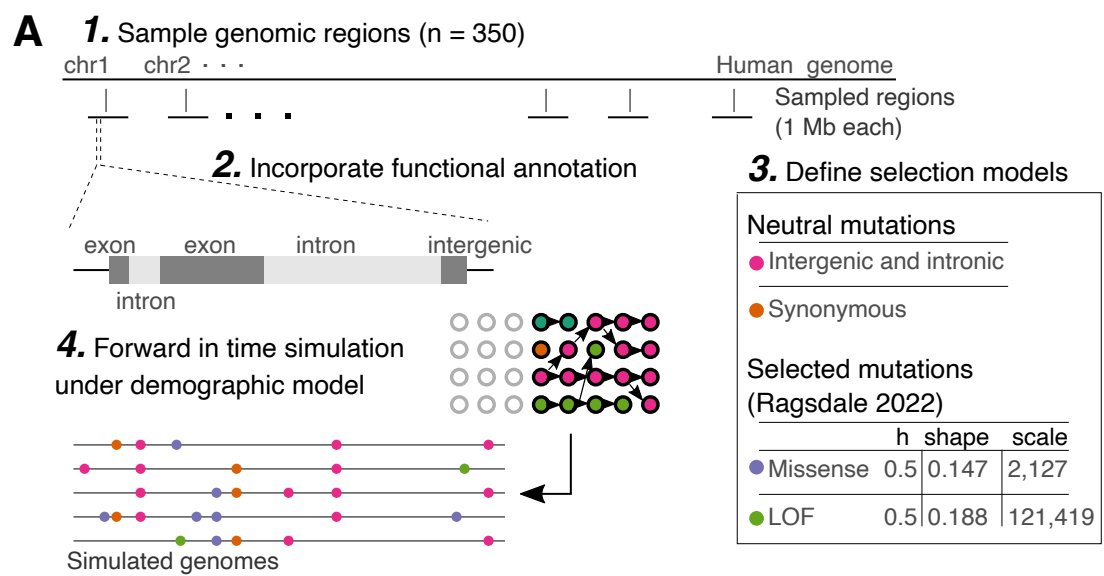

**Figure S6.** Forward in time simulations of functional genetic variation. (A) We first selected 350 random regions of the human genome, each 1Mb in size, and annotated the functional elements within each region, including exons, introns, and intergenic regions. We also calculated the scaled mutation rate for each functional category. Next, we defined the distribution of fitness effects (DFE) for each type of mutation. Finally, we carried out the forward in time simulation under the inferred demography.

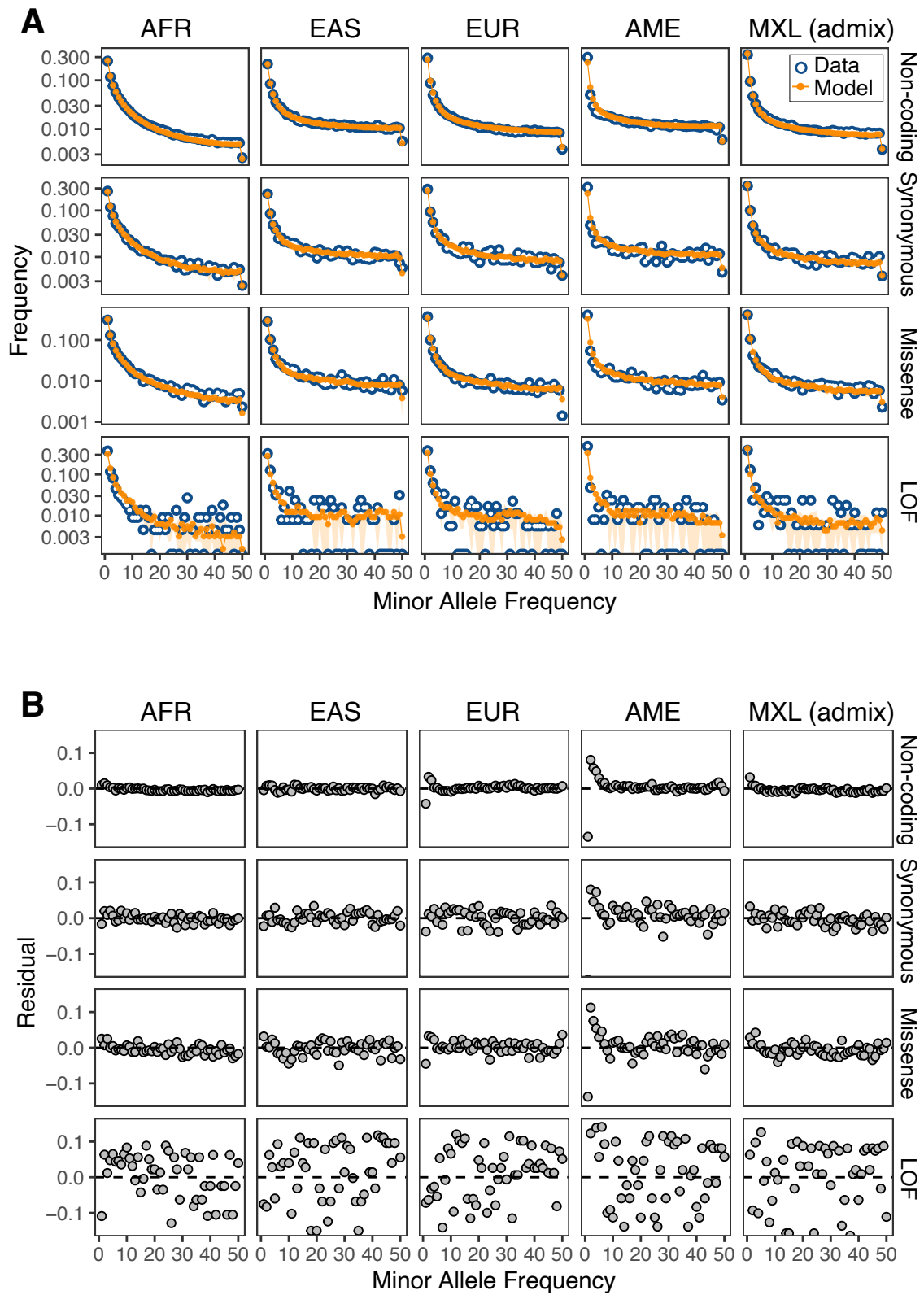

**Figure S7.** Fit of simulated folded Site Frequency Spectrum (SFS) to the Data (A) The resulting comparison, shown as a folded site frequency spectrum, demonstrates a strong match between the observed and simulated data across different functional categories and populations. The blue lines represent the observed data SFS, while the orange lines represent the simulated SFS. Each column represents a different population. (B) Poisson residuals of SFS simulated fit to the data. Overall we observe a good fit between the simulated and observed data. Poisson residuals were calculated as the difference between the observed and expected frequencies, normalized by the square root of the expected frequencies. Overall, we observe a good fit between the simulated and observed data, with Poisson residuals centered around zero. However, the fit is slightly worse in the AME (Indigenous American) population, where the residuals are more positive for low frequency variants. **8/8**
